# Extracellular vesicles from chylomicron-treated endothelial cells drive macrophage inflammation

**DOI:** 10.1101/2025.02.28.640926

**Authors:** Anna Tilp, Dimitrios Nasias, Andrew Carley, Min Young Park, Ashley Mooring, Munichandra Babu Tirumalasetty, Nada A. Abumrad, Yang Wang, Qing Robert Miao, Douglas Lewandowski, José Alemán, Ira J. Goldberg, Ainara G. Cabodevilla

## Abstract

**Background:** Movement of circulating lipids into tissues and arteries requires transfer across the endothelial cell barrier. This process allows the heart to obtain fatty acids (FAs), its chief source of energy and apolipoprotein B (apoB)-containing lipoproteins to cross the arterial endothelial barrier leading to cholesterol accumulation in the subendothelial space. Multiple studies have established elevated postprandial triglyceride-rich lipoproteins (TRLs) as an independent risk factor for cardiovascular disease (CVD). We explored how chylomicrons affect ECs and transfer their FAs across the EC barrier.

**Methods:** We had reported that media from chylomicron-treated ECs leads to lipid droplet (LD) formation in macrophages. To determine the responsible component of this media, we assessed whether removing the extracellular vesicles (EVs) would obviate this effect. EVs from control and treated cells were then characterized by protein, lipid and microRNA (miR) content. We also studied the EV-induced transcription changes in macrophages and ECs and whether knockdown of scavenger receptor-BI (SR-BI) altered these responses. In addition, using chylomicrons labeled with [^13^C]oleate, we studied the uptake and release of this labed by ECs.

**Results:** Chylomicron treatment of ECs led to an inflammatory response that included production of EVs that drove macrophage LD accumulation. The EVs contained little free fatty acids and triglyceride, but abundant phospholipids and diacylglycerols. In concert with this, [^13^]C labeled chylomicron triglycerides exited ECs primarily in phospholipids. EVs from chylomicron treated versus untreated ECs were larger, more abundant, and contained specific miRs. Treatment of macrophages and naïve ECs with media from chylomicron-treated ECs increased expression of inflammatory genes.

**Conclusions:** EC chylomicron metabolism produces EVs that increase macrophage inflammation and create LDs. Media containing these EVs also increases EC inflammation, illustrating an autocrine inflammatory process. FAs within chylomicron triglycerides are converted to phospholipids within EVs. Thus, EC uptake of chylomicrons constitutes an important pathway for vascular inflammation and tissue lipid acquisition.

## INTRODUCTION

How do lipids within lipoproteins leave the circulation and enter tissues, like the heart, as well as the arterial wall? Although these transendothelial movements of lipids are needed for normal physiology, they also are likely responsible for several cardiovascular diseases (CVDs). Atherosclerosis and its associated diseases remain the top cause of morbidity and mortality worldwide ^1,2^. The development of atherosclerosis requires a cascade of events, including chronic inflammation of endothelial cells (ECs), subendothelial lipid accumulation, and the recruitment and infiltration of immune cells with development of lipid-laden macrophages ^3^. Although progression of atherosclerosis is linked to increased circulating levels of LDL ^4^, other lipoproteins that contain apolipoprotein B (apoB) also appear to drive atherosclerosis development. Population studies have shown that measurements of apoB levels or non-HDL cholesterol, cholesterol within LDL and triglyceride (TG) -rich lipoproteins (TRLs), track with CVD better than LDL alone ^5,6^. These data suggest that TRLs cause or exacerbate atherogenesis. TRLs (mostly chylomicrons) are increased in the postprandial period, and greater postprandial lipemia is an independent risk factor for cardiovascular events ^1^.

TRLs are also a source of FAs for the heart; FA oxidation is the primary engine driving heart function. Heart FAs are primarily obtained via lipoprotein lipase (LpL) hydrolysis of TGs. Loss of this pathway leads to smaller hearts^7^ that cannot respond normally to increased afterload due to hypertension^8^ and acute aortic coarctation^7^. Heart failure leads to reduced LpL expression and FA oxidation REF, a process that might be alleviated if FA supplies are maintained. Could there be alternative lipid uptake pathways that could overcome defective lipolysis?.

Emerging studies also point towards the importance of ECs in the initiation and progression of atherosclerosis and CVD ^9^. Dysregulation of EC function marks an early hallmark of atherosclerosis and triggers inflammatory signaling events that are communicated to other cells and contribute to plaque formation (reviewed in ^10^).

Unlike in capillaries that utilize lipoprotein lipase (LpL) to release NEFAs from chylomicrons^11^, the LpL-binding protein glycophosphoinositol HDL binding protein 1 (GPIBHP1)^12^ is not expressed in most arterial ECs^12^. Nonetheless, arterial ECs develop lipid droplets (LDs) during the postprandial period ^13^ due to internalization of unhydrolyzed chylomicrons via scavenger receptor-BI (SR-BI)^14^. Internalized chylomicrons are subsequently hydrolyzed in the lysosomal compartment, a process that leads to EC LD accumulation. Chylomicron uptake by ECs also leads to LD accumulation in skin macrophages of hyperchylomicronemic, LpL-deficient mice^14^. The chylomicron-treated ECs also produce media that promote LD accumulation by peritoneal macrophages (PMACs)^14^. This lipogenic effect on PMACs was ablated upon EC knockdown of SR-BI or inhibition of lysosomal activity in chylomicron treated ECs, indicating that both uptake and lysosomal degradation are involved in this process^14^.

We hypothesized that chylomicron treatment leads to EC lipid transfer from the apical to the basolateral side. This process might allow transendothelial transfer of chylomicron derived FAs but also alter the biology of ECs. Chylomicrons undergo lysosomal hydrolysis within ECs ^14,15^ and could allow release of either intact non-polar lipids or lipids hydrolyzed from the EC basolateral surface. We recently reported that NEFAs associated with CD36 induce creation and secretion of small EVs ^16^ and others showed that ECs are a major cellular source of EVs ^17–20^. EVs may transmit biological information that alter PMACs due to their cargo of nucleic acids, proteins and lipids, and have been previously correlated with atherosclerosis ^21^.

In this report, we show that chylomicron uptake induces expression of inflammatory genes in ECs. In addition, we demonstrate that EV release by ECs is required for development of LDs in co-cultured macrophages. The EVs are devoid of TGs and primarily composed of phospholipids and diglycerides with the FAs from chylomicron TGs exiting ECs in EV phospholipids. Chylomicron treatment increases EC-derived EV number and their associated miRs. In addition to LD accumulation, these EVs induce an inflammatory gene expression profile in macrophages and other ECs. Our data suggest that EC chylomicron metabolism provides a pathway for lipid delivery to subendothelial cells and triggers macrophage and endothelial inflammation.

## METHODS

### Cells

All cells were cultured and maintained according to established guidelines and manufacturés statements in a standard humidified incubator at 37°C and 5% CO_2_. For experiments, cells were seeded in respective gelatin pre-coated (Millipore Sigma; EmbryoMax 0.1% Gelatin Solution) cell-culture dishes or glass coverslips for microscopy.

### Murine Endothelial Cells (MECs)

MECs (Angiocrine Bioscience) were cultured in Advanced DMEM/Ham’s F-12, supplemented with 20% FCS, 1% penicillin/streptomycin, 10 mM HEPES buffer, 1% GlutaMax, 5 µM SB 431542, 50 µg/ml heparin, 20 ng/ml FGF, 10 ng/ml VEGF, and 50 µg/ml BT-203 endothelial mitogen (full medium). Starvation medium contained all supplements excluding FBS. For the period of EV production, cells were kept in minimum medium, without FBS, FGF and VEGF.

### Primary Murine Peritoneal Macrophages (PMACs)

PMACs were isolated from 6-week-old wild type (WT) mice as previously described [10]. The animal procedures were approved by the Institutional Animal Care and Use Committees at New York University Langone Health Medical Center. Briefly, mice were anesthetized in isoflurane chamber and subsequently euthanized by cervical dislocation. Next, they were injected with 4 ml ice-cold PBS into the peritoneal cavity. After a gentle massage of the abdomen, at least 80% of injected PBS was collected through a small skin incision. Cells were pelleted by centrifugation (300g for 10 minutes at 4°C) and seeded in RPMI medium supplemented with 10% FCS and 1% penicillin/streptomycin in transwell culture plates to allow PMACs to attach overnight. After 24h, plated cells were washed with PBS to eliminate non-adherent cells.

### SR-BI ASO Treatment

Knockdown of SR-BI in MECs was performed one day before FCS starvation using ASO transfection with DharmaFECT4 (Fisher Scientific) transfection reagent according to manufactureŕs statement. SRBI ASO (GCTTCAGTCATGACTTCCTT, ISIS no. 353382) and control were provided by Ionis Pharmaceutical.

### Chylomicron Treatment

MECs were treated with chylomicrons as described previously [10]. Briefly, cells were seeded in 20cm^2^ cell culture dishes in complete medium. Before chylomicron treatment, cells were deprived of FCS for at least 16h. Chylomicrons were applied to cells in a concentration of 4 mg/dL TG in FCS free medium for 30 minutes. After thoroughly washing with PBS, medium was changed to minimum medium and was collected 24h after chylomicron treatment. EV-containing medium was either used as conditioned medium or for isolation of EVs.

### Isolation of Small Extracellular Vesicles

Culture medium harvested from ECs after chylomicron pulse or control conditions was centrifuged at 1000xg for 10 minutes to remove cell debris and concentrated using filter units (Amicon). EVs were isolated using the miRCURY Exosome Kit for cell culture (Quiagen) according to manufactureŕs directions. Vesicles were washed with PBS and re-suspended directly in lysis buffer for miR isolation (miRNeasy Micro Kit, Quiagen) or PBS for other experiments.

### Treatment with Conditioned Medium

Conditioned medium from chylomicron-treated and control cells was harvested after 24h. Medium was centrifuged at 1000xg for 10 minutes to remove cell debris. PMACs and naïve ECs were seeded in pre-coated 6 well plates and treated with conditioned medium for 24h.

### RNA Isolation from Cells

RNA was isolated from cells after thoroughly washing with PBS, PMACs were directly harvested in lysis buffer (RNeasy Micro Kit, Quiagen) and RNA was isolated according to manufactureŕs statement. Naïve ECs were harvested directly in TRIzol reagent and RNA was isolated using chloroform-phenol phase separation. After a wash with 75% ethanol, RNA was dissolved in RNAse free water and subjected to RNA Sequencing.

### RNA Sequencing and Pathway Analysis

Total RNA was extracted from cultured MECs and PMACs as was described in the previous section. NYU Langone’s core genomic facility performed quality control analysis of RNA samples, library preparation and subsequent sequencing analysis. RNA sequencing data was generated from one lane of a paired-end 50 Illumina NovaSeq X 10B run. Next, per-read per-sample FASTQ files were generated using the bcl2fastq2 Conversion software (v2.20) to convert per-cycle BCL base call files outputted by the sequencing instrument into the FASTQ format. The alignment program, STAR (v2.7.3a), was used for mapping reads of every sample to the mouse reference genome mm10. The feature Counts (Subread package v1.6.3) software was used to generate matrices of read counts for annotated genomic features. Differential expression statistical analysis was performed by the DESeq2 package in the R programming environment (R v4.1.2) to compare the group of PMACs treated with condition medium from chylomicron-treated ECs and control conditions. Similar comparison was performed from the EC dataset. Differentially expressed genes were annotated into Kyoto Encyclopedia of Genes and Genomes (KEGG) pathways by applying a cut-off for the adjusted p-value (q<0.05). Significant results were visualized in the R statistical environment (R v4).

### miR Sequencing

The analysis of count data generated from RNA-seq was used to detect the differentially expressed genes. The package DESeq2 through R/Bioconductor provides a method to test for differential expression by use of negative binomial generalized linear models. The starting point of a DESeq2 analysis is a count matrix K with one row for each gene i and one column for each sample j. The matrix entries K ij indicate the number of sequencing reads that have been unambiguously mapped to a gene in a sample. The filter statistic in DESeq2 is the mean of normalized counts for a gene, while the test statistic is p, the P value from the Wald test. We considered significant all genes with a fold change greater 0.5 or lower than 0.5 and corrected P value less than 0.5 (FDR Test). To gain more mechanistic insights into the underlying biology of the differentially expressed genes, we employed (KEGG) pathway analysis. Statistically significant related pathways were filtered through g:profiler tool with a P value less than 0.5 for each category.

### Collection and Culture of PMACs

WT C57BL/6J mice were given an intraperitoneal injection of zymosan (0.1 mg in 200 μL of PBS) to induce mild inflammation. At 72 hours after the injection, mice were anesthetized by isoflurane inhalation and euthanized by cervical dislocation. An incision was performed to expose the peritoneal membrane, and 5 mL of ice-cold PBS was injected into the peritoneal cavity, followed by a gentle massage of the abdomen. Using surgical scissors, an incision was performed in the peritoneal membrane, and at least 4.5 mL of liquid was collected with a 1000 μL pipette into a 15 mL conical tube. PMACs were pelleted by centrifugation (300g for 10 minutes at 4°C) and re-suspended in 1mL RPMI 1640 culture medium (ThermoFisher Scientific). PMACs were plated in glass coverslips at 0.2 × 106 cells/mL in RPMI 1640 culture medium supplemented with 10% FBS, 1% penicillin/streptomycin (pen/strep), 10 μM sodium pyruvate, 25 mM HEPES, and 100 μM amino acids. PMACs were maintained at 37°C and 5% CO_2_ in a humidified incubator. All experiments were performed within 48 hours after PMAC collection.

### Treatment of PMACs with EC Conditioned Media

Cultured MECs were either left untreated (control) or pulsed with chylomicrons for 30 minutes, washed thoroughly, and switched to FBS-free culture medium for 24 hours. Conditioned media were collected and EVs isolated as described above. Cultured PMACs were treated for 16 hours with complete (EV-containing) conditioned media from control or chylomicron-pulsed MECs, EV-free media from chylomicron-pulsed MECs, or control media containing EVs isolated from chylomicron-pulsed MEC conditioned media. Following treatment, PMACs were washed twice with PBS, then fixed for 20 minutes with formalin at RT. LD staining with BODIPY 493/503 was performed by incubation with 1:1000 by volume of a BODIPY 493/503 stock solution (5 mM in DMSO) for 30 minutes at RT. Nuclei were highlighted with DAPI. Samples were imaged with a Leica SP8 confocal microscope. LDs were quantified from microphotographs using ImageJ/FIJI.

### Single Particle Interferometric Reflectance Imaging Sensor (SP-IRIS) Analysis

Tetraspanin CD9, CD63 and CD81 distribution were analyzed using the ExoViewR100 platform using the Leprechaun mouse tetraspanin ExoView kits (Unchained Labs Catalog number 251-1046) following the kit assay protocol. Briefly, fraction 2 from the density gradient was diluted 4 times with the incubation solution II. Fifty μL of the dilute samples were placed on top of the chips inside a Falcon 24-well cell culture plate, flat bottom (Fisher Scientific Catalogue number 08-772-1) for the capture of EV carrying rat anti mouse CD9 (Clone MZ3), hamster anti mouse CD81antigen (Clone Eat-2) and Armenian hamster isotype IgG (Clone HTK888) and rat isotype IgG2ak (Clone RTK2758). Wells were sealed with adhesive plate seals to prevent evaporation. After 16 h incubation at room temperature, chips were washed 3 times with 1 X solution A for 3 minutes on ELISA microplate orbital shaker at 500 rpm (FisherbrandTM Fisher Scientific Catalogue number 88-861-023). Chips were then incubated with an antibody cocktail made of 0.6μL Armenia hamster anti mouse CD81+(Clone Eat-2) conjugated with AlexaFluor 555, 0.6μL rat anti mouse CD63(Clone NVG-2) conjugated with AlexaFluor 647, and 0.6 μL of rat anti mouse CD9 (Clone MZ3) conjugated with AlexaFluor 488 in 300μL of blocking solution for 1h at RT in on orbital shaker at 500 rpm. Chips were washed 1 time with 1X solution A, 3 times with 1X resolution B and 1 time with DI water respectively for 3 minutes at 500 rpm. After drying the chips, image acquisition from each chip was carried out using the ExoView® R100 platform, and the data were analyzed by the ExoView Analyzer software version 3.2 (NanoView Biosciences). The images of the acquisition were visually inspected and all the artifacts onto the spots were manually removed from the analysis. Non-specific binding was checked on isotype controls IgG spots. The cut off was manually established for all the chip to exclude the majority of the signal (> 90%) captured on the isotype control.

### Production of ^13^C-labeled Chylomicrons

Micelles were prepared by adding 6 mmol/L oleate (U-^13^C_18_), 2.5 mmol/L 1-oleoyl-rac-glycerol, 1.5 mmol/L 1-palmitoyl-2-hydroxy-sn-glycero-3-phosphocholine, and 0.6 mmol/L L-α phosphatidylcholine to 10 mL of chloroform in a glass tube. The chloroform was evaporated under N_2_ gas and 20 mL of PBS containing 12 mmol/L sodium taurocholate was added to the tube and vortexed for 1 min. The lipid suspension was sonicated until optically clear. The micelle solution was used for rat duodenum infusion and lymph collection within 12 hrs.Thoracic lymph duct cannulation and lymph collection was performed in male Sprague-Dawley rats (340–480 g) (Jackson Laboratories) as described previously, with modifications ^22^. Rats were fasted overnight and anesthetized with etomidate (10 mg/kg) and isoflurane prior to intubation after which anesthesia depth was maintained by 1-2 % isoflurane along with the analgesic carprofen (5 mg/kg). The abdomen was opened via a 3 cm left subcostal incision, and the thoracic duct was cannulated with saline-rinsed, pre-cut catheter (16 gauge) connected to a collection tube. The micelle solution was infused into the duodenum at a rate of 3 ml/min (Harvard pump) followed by PBS. The abdomen was closed and lymph was collected for 6-8 hrs. Body temperature was maintained at 37°C via a heating pad. The TG content of collected fractions were analyzed by LC-MS, as described previously ^23^. The lymph collected from two rats was combined and filtered through a 5 µm syringe filter. ApoCII (50 ug) was added to filtered lymph and the chylomicrons were stored at 2-8°C. These procedures involving rats were approved by the Institutional Animal Care and Use Committees (IACUC) at the Ohio State University.

### Targeted Lipidomic Analysis by LC-MS

Exosome and chylomicron samples were deproteinated with cold methanol prior to lipid extraction, and Avanti Splash lipidomix^®^ (Birmingham, AL) was added to each sample as an internal standard (Splash:sample, 1:94). The non-polar lipids and polar metabolites were extracted using a modified Bligh-Dyer chloroform/methanol/water extraction method. Polar metabolites and non-polar metabolites were separated into two layers post centrifugation. The top layer is a polar water/methanol layer, while the bottom layer is nonpolar chloroform. The non-polar chloroform layer was centrifuged in vacuum at 40°C and re-suspended in 40uL of a 4:3:1 (v/v) isopropanol/acetonitrile/water solution. Analysis of nonpolar fractions was performed using an Agilent Infinity II 1290 liquid chromatography system coupled to an Agilent 6230B Time-of-flight mass spectrometry platform (Santa Clara, CA). Metabolite and lipid separation during chromatography was performed through Acquity Premier CSH C18 column (130Am 1.7μm, 2.1×150mm). Column temperature was programmed at 50°C with a biphasic gradient change in mobile phase using a 5:1:4 solution of isopropanol/methanol/water and 5mM ammonium acetate and 0.1% acetic acid buffers on pump A and a solution of 99:1 (v/v) of isopropanol/water and 5mM ammonium acetate and 0.1% acetic acid buffer on pump B. Lipid profiles were analyzed using the proprietary Agilent platform MassHunter and Mass Profiler Pro (Santa Clara, CA) against the Metlin database. For [^13^C]oleate labelling experiment, identified lipid compounds containing 18:1 were further selected with area threshold >5000 and quantified for M+0, M+18, M+36, M+54 with as a measure of [^13^C]oleate incorporation. Labeled lipid profile was calculated as the percentage of labeled family or classes in the sample. Graphing was performed in MassProfiler Pro and GraphPad Prism.

### Tracking [^13^C]oleate in Endothelial Cells and Extracellular Vesicles

Lipidomic analysis and [^13^C}oleate content of ECs and conditioned media were performed via LC-MS/MS at NYU Metabolomics Core. Lipid extraction of the samples was conducted as previously ^24^. Lipidomic data were obtained by putatively identifying peak heights using LipidBlast spectral library for named lipids top scoring hits (RevDot>900). Untargeted features were selected using similar methodology, and all named lipids and lipid features were processed using down-stream statistical pipelines written by the NYU Metabolomics Core Resource Laboratory. The resulting ThermoTM RAW files were converted to SQLite format using an in-house python script to enable downstream peak detection and quantification. The available MS/MS spectra were first searched against the LipidBlast Spectral Library and respective Decoy spectral library databases using an in-house data analysis python script adapted from our previously described approach for metabolite identification false discovery rate control (FDR) ^25,26^. Putatively identified metabolites were filtered for any duplicated metabolites names to generate a list of metabolites with unique names. Next, the decoy hits in the resulting list were dropped, and the peak heights for each putative metabolite hit were extracted from the sqlite3 files based on the metabolite retention time ranges and accurate masses in the above-mentioned metabolite list. For isotope labeling analyses, we calculated the theoretical m/z value for each isotopologue based on the metabolite formula, ion type, and combination of isotopes, including no labeling through maximum labeling. These metabolite peaks were then extracted based on the theoretical m/z of the expected ion type (with or without labeling), e.g., [M+H]+, with a 15 part-per-million (ppm) tolerance and a ± 0.2 min peak apex retention time tolerance within an initial retention time search window of ±0.5 min. The resulting data matrix of metabolite intensities for all the samples and blank controls was processed using an in-house python script (https://github.com/NYUMetabolomics/plz), and the final peak detection was calculated based on a signal-to-noise ratio (S/N) of 3× compared to the blank controls, with a floor of 10,000 (arbitrary units). For the samples where the peak intensity was lower than the blank threshold, the metabolites were annotated as not detected and were imputed with either the blank threshold intensity for statistical comparisons so as to enable an estimate of the fold change, as applicable, or zeros for the median metabolite intensity calculation of a sample. For isotope labeling analyses, we further converted the resulting peak intensities to relative abundance to the monoisotopic peak (defined as 100%) for each metabolite and converted to a percentage value. Unlabeled control samples were used to assess isotope labeling enrichment and background levels of natural isotopes. Peak heights were analyzed using Xcalibur (ThermoFisher Scientific) based on previously established Lipidblast library. For ^13^C-oleate labelling experiment, fatty acids and TGs were selected and quantified for M+0, M+18, M+36, M+54 with as a measure of [^13^C]oleate incorporation. Labeled lipid profile was calculated as the percentage of labeled family or classes in the sample. Graphing was performed in GraphPad Prism.

## RESULTS

### Chylomicron treatment induces expression of inflammatory genes in ECs

We had shown that ECs internalize undigested chylomicrons via a pathway involving SR-BI ^14^. Here, we assessed how this uptake affects ECs by measuring EC gene expression before and after chylomicron exposure. Chylomicron treatment resulted in transcriptional remodeling of ECs (**Fig. 1A** and **Supplement Table 1**). The most highly induced genes were involved in innate immunity processes, e.g. chemokines (e.g. *Cxcl2, Cxcl5*), cytokines (e.g. *Ccl2, Ccl7*), and complement system (e.g. *C3*), or were key mediators of inflammation (e.g. *Saa3, Lcn2*). Among the highly induced genes were *Vcam1* and *Icam1* (**Fig. 1 A** and **B**), which play a critical role in leukocyte-endothelial interaction in inflammation ^27,28^. Others have shown that EC LD accumulation due to defective TG lipolysis leads to a similar increase in these genes in the aorta ^29,30^. To better understand biological relevance of the differentially expressed genes, we performed KEGG pathway statistical analysis **(Fig. 1C)**. The most enriched categories included pathways of inflammation and immunity, such as “TNFα signaling”, “NF-kappa B signaling”, and “IL-17 signaling”, or categories that link these genes directly to CVD, such as “lipid and atherosclerosis” (**Fig. 1C, Supplement Table 1**). Expression values comparing genes enriched in these pathways for the chylomicron-treated versus untreated ECs are shown in heatmaps (**Fig. 1D**). Hence, exposure of ECs to chylomicrons *in vitro* induces an inflammatory response with gene signatures involved in vascular dysfunction and cardiovascular disease.

**Figure 1.**
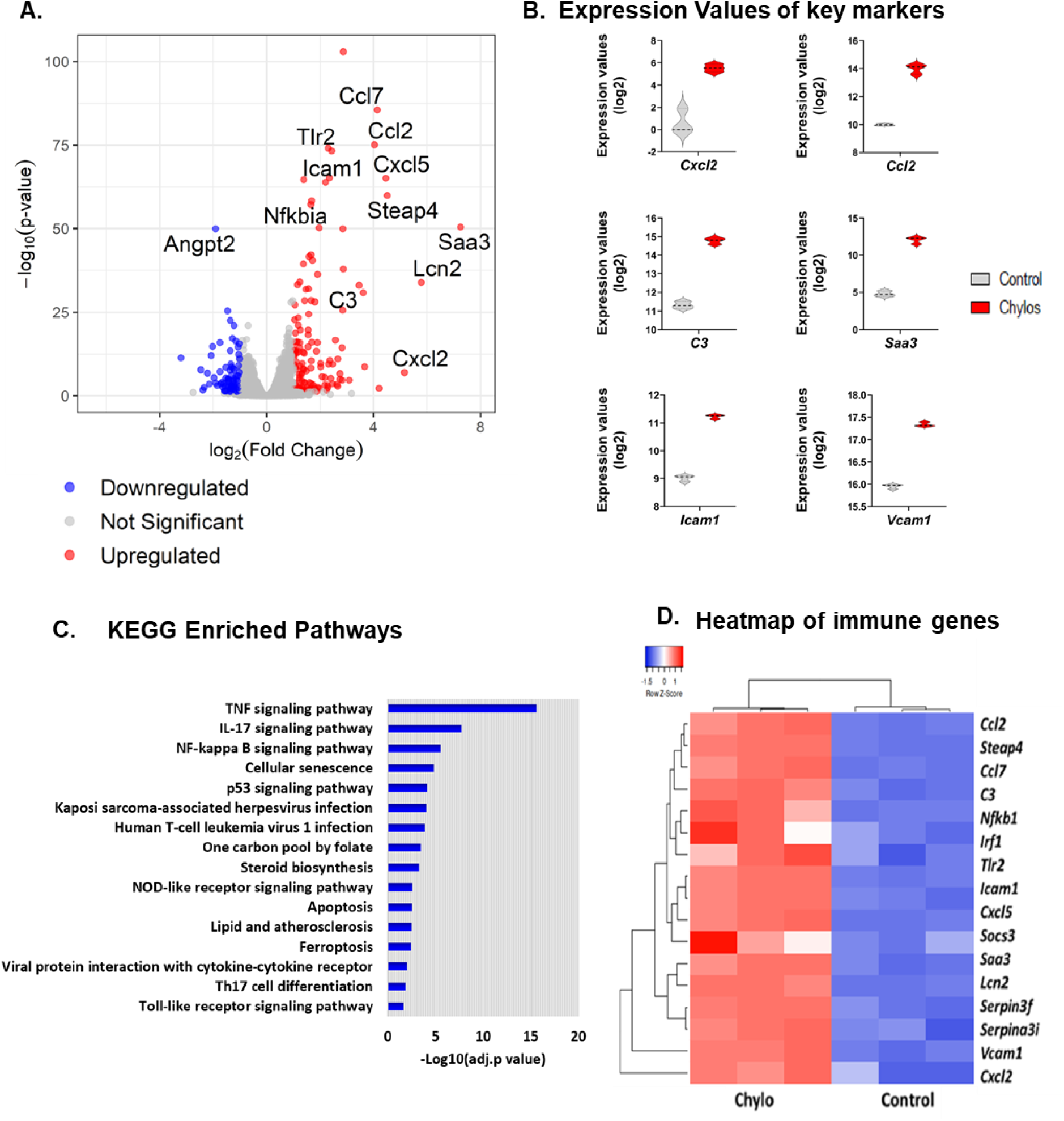
Differential expression analysis of chylomicron-treated versus control endothelial cells (EC). (**A**) Volcano plot illustrating significantly altered gene expression, determined by DESeq2 analysis and threshold values (absolute log2 fold change > 1 and adjusted p < 0.05). Dashed lines and shaded regions indicate these thresholds, with red showing upregulated genes, blue indicating downregulated genes, and gray showing non-significant changes. Adjusted p-values are shown on a –log10 scale. (**B**) Violin plots of log2-transformed expression levels for representative inflammatory and adhesion-related genes in control (gray) versus chylomicron-treated (red) cells. Data represents three independent biological replicates (n = 3). (**C**) KEGG pathway enrichment analysis (adjusted p < 0.05) highlighting immune-related pathways among the upregulated genes. KEGG pathways were ranked by statistical significance, with the bar length representing the adjusted p-value score on a –log10 scale. Longer bars indicate higher significance. (**D**) Heatmap of genes enriched in immune-related KEGG pathways (adjusted p < 0.05). Hierarchical clustering was performed using Euclidean distance and average linkage. Colors represent z-scored expression values, with red indicating higher expression and blue indicating lower expression.

### EC-derived EVs induce LD accumulation in macrophages

We previously reported that EC uptake and lysosomal hydrolysis of chylomicrons triggers LD accumulation in co-cultured macrophages ^14^.To our surprise, this effect was not due to EC release of NEFAs following lysosomal chylomicron hydrolysis, as assessed by ultra-performance liquid chromatography coupled with quadrupole time-of-flight mass spectrometry (UPLC-TOF-MS)^14^. Electron microscopy analysis of chylomicron-treated EC conditioned media (CEM) revealed the presence of EVs (**Fig. S1A**). To investigate whether EC-derived EVs are involved in the induction of LDs in macrophages, we treated PMACs with conditioned medium from control or chylomicron treated ECs. A schematic of this experiment is shown in **Fig. 2A**. Briefly, ECs deprived of FBS overnight, were left untreated (control) or pulsed with chylomicrons (4mg/dL TG) for 30 minutes in FBS-free medium. Following treatment, ECs were washed thoroughly with PBS and maintained in FBS-free medium for 24hs. Conditioned media were collected and a subset filtered using MilliporeSigma™ Amicon™ Ultra-15 Centrifugal Filter Units. This allowed for generation of EV-containing and EV-free conditioned medium, as well as a concentrated fraction of EC-released EVs.

**Figure 2.**
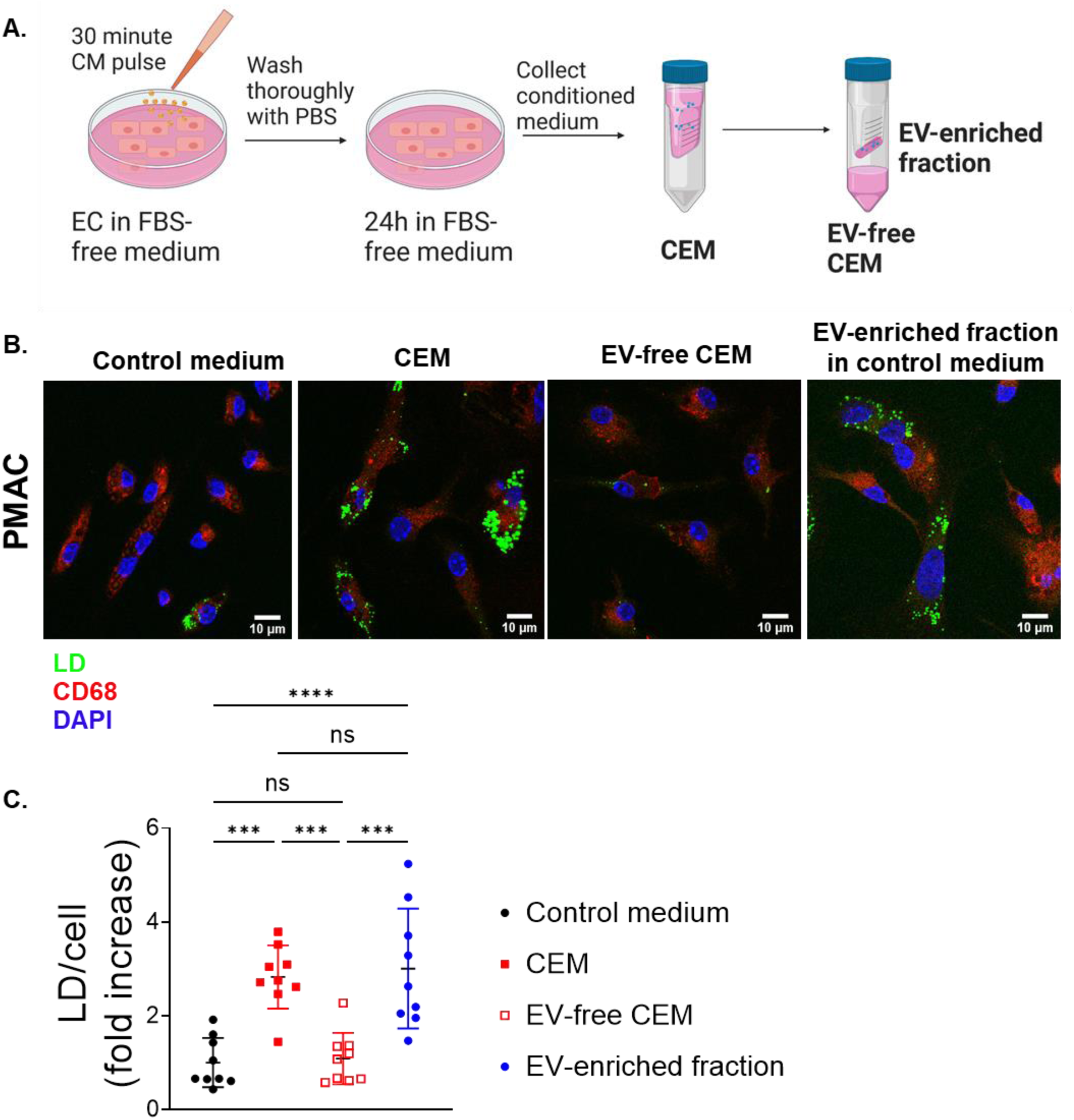
EC-derived small extracellular vesicles are responsible for LD accumulation in cultured primary macrophages. **A**. Experimental overview of conditioned medium production. Murine endothelial cells are cultivated in normal expansion medium and deprived of FBS at least 24h prior to chylomicron treatment. Chylomicrons (4 mg/dl TG) are applied for 30 minutes and medium was changed after thoroughly washing with PBS. Conditioned medium of untreated (control) cells and chylomicron pulsed cells was harvested after 24h. EVs were removed from medium through centrifugation with filter units. **B**. Representative microscopy images of macrophages treated with indicated conditioned medium from ECs. Macrophages treated with medium from chylomicron treated ECs display increased lipid droplet accumulation compared to cells treated with control conditioned medium or after removal of EVs. Cells were stained with DAPI (blue), CD68 (red) and BODIPY (green) and imaged using confocal microscopy. Quantification of 9 independent experiments shown in **C**. ***p <0.0001, ****p<0.00001, ANOVA Multiple comparisons test.

As observed in previous co-culturing experiments [10], treatment with EV-containing CEM (but not with conditioned media from control, untreated ECs) resulted in LD accumulation in macrophages. Macrophages treated with EV-free CEM did not accumulate LDs, but supplementation of control medium with the EV-enriched fraction obtained from CEM led to PMAC LD formation (**Fig. 2B**). Similarly, naïve ECs treated with the same conditioned media developed LDs (**Fig. S1B**). These results indicate that endothelial EVs drive LD accumulation in both macrophages and naïve ECs.

### Characterization of EC-released EVs

We next assessed whether chylomicron treatment affected the number, size and surface markers of EC-derived EVs using ExoView R100 chip arrays. The particles were analyzed upon capture with the characteristic EV tetraspanin proteins CD81 and CD9. As shown in **Fig. 3A**, chylomicron treatment of ECs increased 24-hour release of both CD81 and CD9 captured EVs. The distribution of surface marker proteins on EVs varies depending on the cellular origin, culturing conditions as well as environmental stimuli ^31^. Furthermore, these changes associate with change in activity, transport and function of these vesicles ^32,33^. However, besides the increase in particle number, the surface distribution of tetraspanins in the EVs released from chylomicron treated ECs was comparable to control cells (**Fig. 3B**), with CD81 showing the highest expression followed by CD9 and CD63. We also evaluated colocalization of the the two tetraspanins. Vesicles from control and chylomicron treated ECs showed a similar CD81 and CD9 colocalization in about one third of the particles upon CD81 (control 36.0% and chylomicron 35.0%) or CD9 capture (control 29.3% and chylomicron 28.2% (**Fig. 3C**). These results slightly differ from tetraspanin composition of vesicles isolated from human ECs, where tetraspanins colocalize in more than 30% of vesicles^33^. In our vesicles this was only true for 5-7% of particles. Together, our results suggest that chylomicron treatment of ECs does not alter the distribution of surface marker proteins but results in increased number of extracellular vesicles with slightly but significantly increased diameter (**Fig. 3D**).

**Figure 3.**
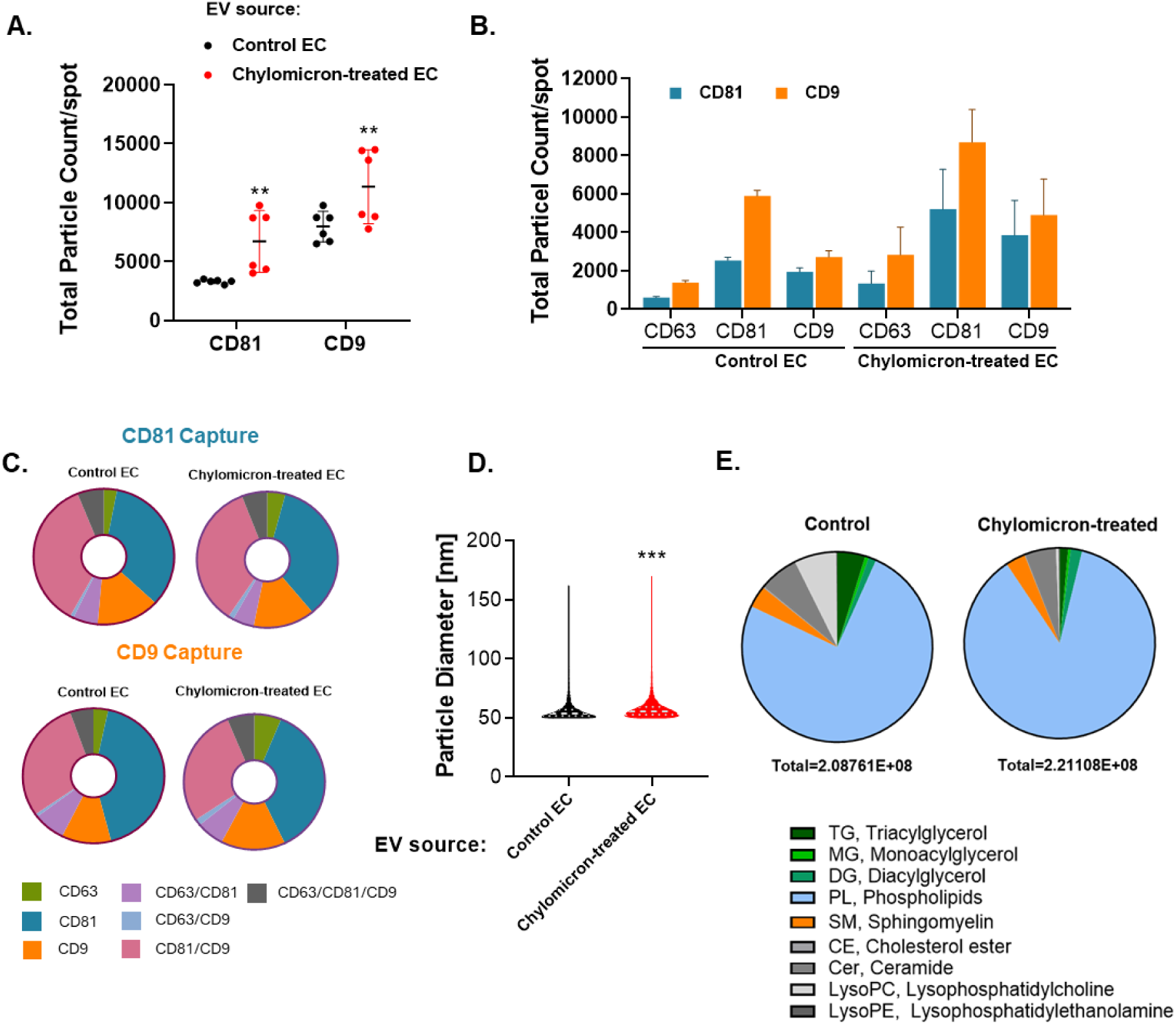
Characterization of EC-released vesicles. Analysis is based on tetraspan detection using the ExoView® R100 platform. **A**. Total number of vesicles is increasing after chylomicron treatment of parent ECs, both on CD81 antibody capture and CD9 capture whereas the distribution of tetraspanins upon respective capture is comparable between the two groups (**B**). **C**. Colocalization [%] of tetraspanins upon CD81 and CD9 capture. Results are represented as average (± SD) from two biological samples and three analyzed spots per sample. **D.** Size Distribution of all particles detected on CD81 capture spot. E. Lipidomics analysis of EV lipid composition.

Lipid composition has been shown to contribute to structure, signaling and uptake of EVs (reviewed in^34^). In general, studies showed an enrichment of cholesterol, sphingomyelin and phospholipids in vesicles from different origins ^32,35,36^. However, information on lipid composition of EC-derived EVs is limited^17^. The lipid composition of the EVs from control or chylomicron treated ECs is shown in **Fig. 3E**. As expected, EVs from both control and chylomicron-treated ECs were enriched in phospholipids, whereas non-polar lipids, such as TGs, were present in only minor amounts.

### Uptake and release of ^13^C-labeled chylomicrons

To determine how chylomicron lipids were processed and possibly re-secreted by ECs, we treated ECs with [^13^C] oleate labeled chylomicrons from rat lymph. A schematic of this experiment is shown in **Fig. 4A**. ECs were either left untreated (control) or pulsed for 30 minutes with [^13^C] oleate labeled chylomicrons (4mg/dL chylomicron TG), then thoroughly washed with PBS, switched to FBS-free medium for 24 hours, after which ECs and conditioned media were collected and processed for lipidomic analysis. As expected, the majority (84.1%) of the [^13^C]oleate label in rat lymph chylomicrons was detected in the TG pool **(Fig. 4B**). In ECs, 13% of the [^13^C] oleate label was detected in the TG pool, which is in keeping with our previous observations that internalized chylomicron lipids fuel LD biogenesis in ECs (10). However, the majority of [^13^C]oleate label was found in the phospholipid pool; 51% in ECs and 95% in the CEM (**Fig. 4C**). The number of [^13^C] oleate moieties in each lipid family differed between ECs and CEM. In chylomicrons, labeled TG contained one [^13^C]oleate moiety (**Fig. 4B**). Intracellular labeled TG was composed of one, two or three [^13^C]oleate moieties at 40.6%, 52% and 7.4%, respectively, whereas all labeled TG in conditioned medium contained one [^13^C]oleoyl side chain (**Fig. 4C**). Similarly, labeled phospholipids (PLs) and sphingomyelins (SMs) in ECs were incorporated with one or two [^13^C]oleate, but majority of labeled PL and SM in conditioned media contained one [^13^C] oleoyl side chain. Moreover, labeled lipid species were comprised of various fatty acid lengths, including the ones with carbon length < C18, e.g. TG(16:1/16:1/16:1). These data indicate that chylomicron TG-bound [^13^C]oleate was intracellularly hydrolyzed and incorporated into TG for LD formation and in PL for the release of EVs. This is in keeping with our previous finding that ECs synthesize LDs following chylomicron uptake and lysosomal hydrolysis ^14^.

**Figure 4.**
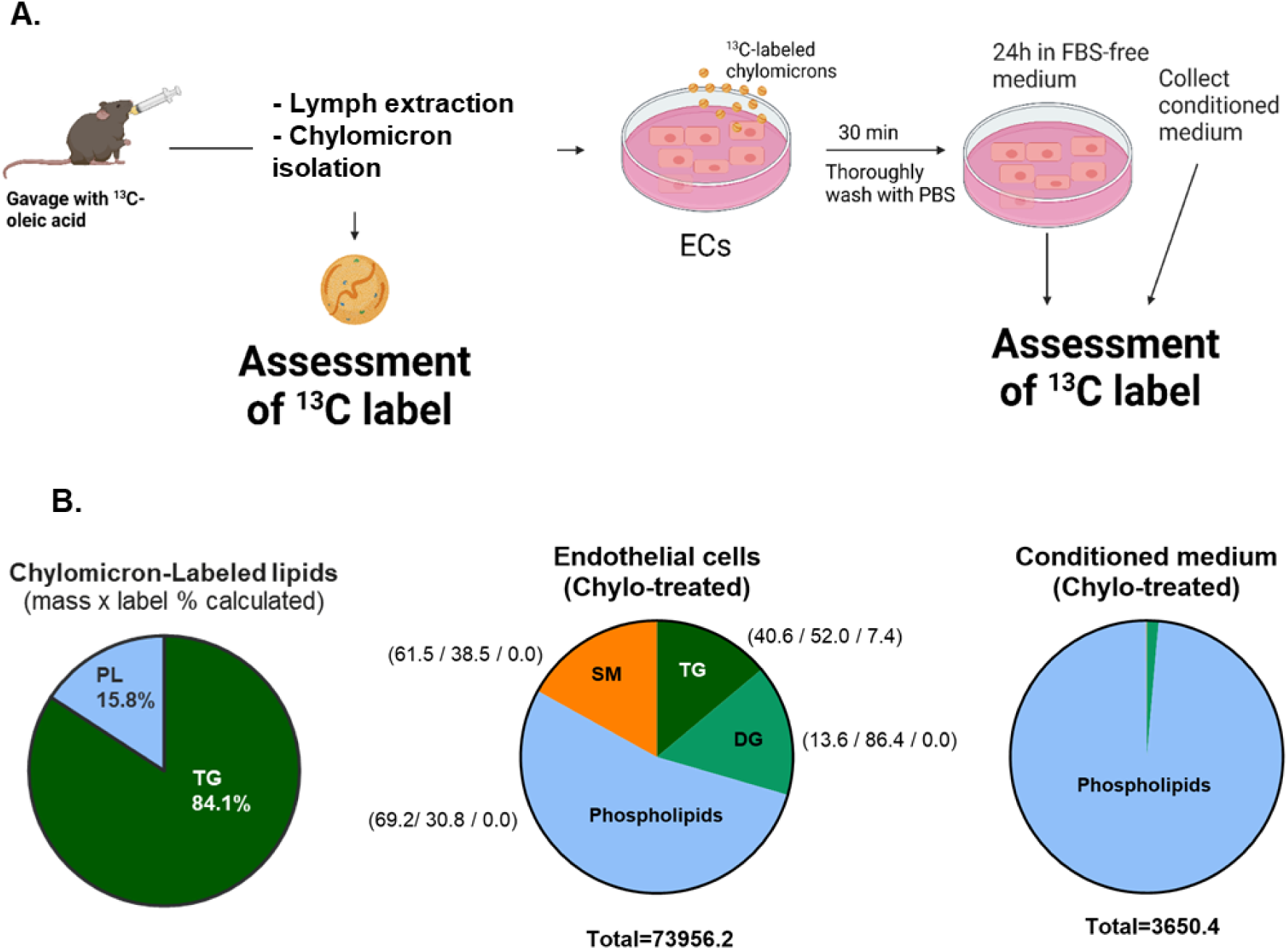
Uptake and release of ^13^C-labeled chylomicrons. **A**. Schematic diagram of [^13^C]oleic acid (OA) chylomicron experiment. **B**. Distribution of [^13^C]OA in chylomicrons. C. Percentage of total [^13^C]OA in endothelial cells and conditioned medium. D. Relative [^13^C]OA in each lipid family or species in endothelial cells and conditioned medium. Values in parenthesis indicate the percentage of 1, 2, or 3 [^13^C] OA moiety in each lipid family or species. TG, triacylglycerol; DG, diacylglycerol; PL, phospholipids; SM, sphingomyelin; EC, endothelial cells; CM, conditioned medium.

### CEM induces inflammatory gene expression in PMACs

LD accumulation in macrophages is a hallmark of inflammatory activation ^37,38^. To determine how CEM affected macrophage biology, we performed RNA sequencing analysis of macrophages treated for 16 hours with conditioned media from control or chylomicron-treated ECs. Treatment with CEM triggered transcriptional remodeling in macrophages (**Fig. 5A, S.Table 1**). Global functional analysis of the differentially expressed genes with KEGG pathways revealed that most of the up-regulated genes were enriched in categories related to innate immunity and inflammation (**Fig. 5B, S. Table 2**). These pathways include major inflammatory cascades such as cytokine-cytokine receptor interaction, TNF signaling and NFkB signaling. We specifically found greater expression of key pro-inflammatory genes such as *Il6, Il1b, NFKb, Vcam1* and also markers of macrophage activation e.g. CD69.

**Figure 5.**
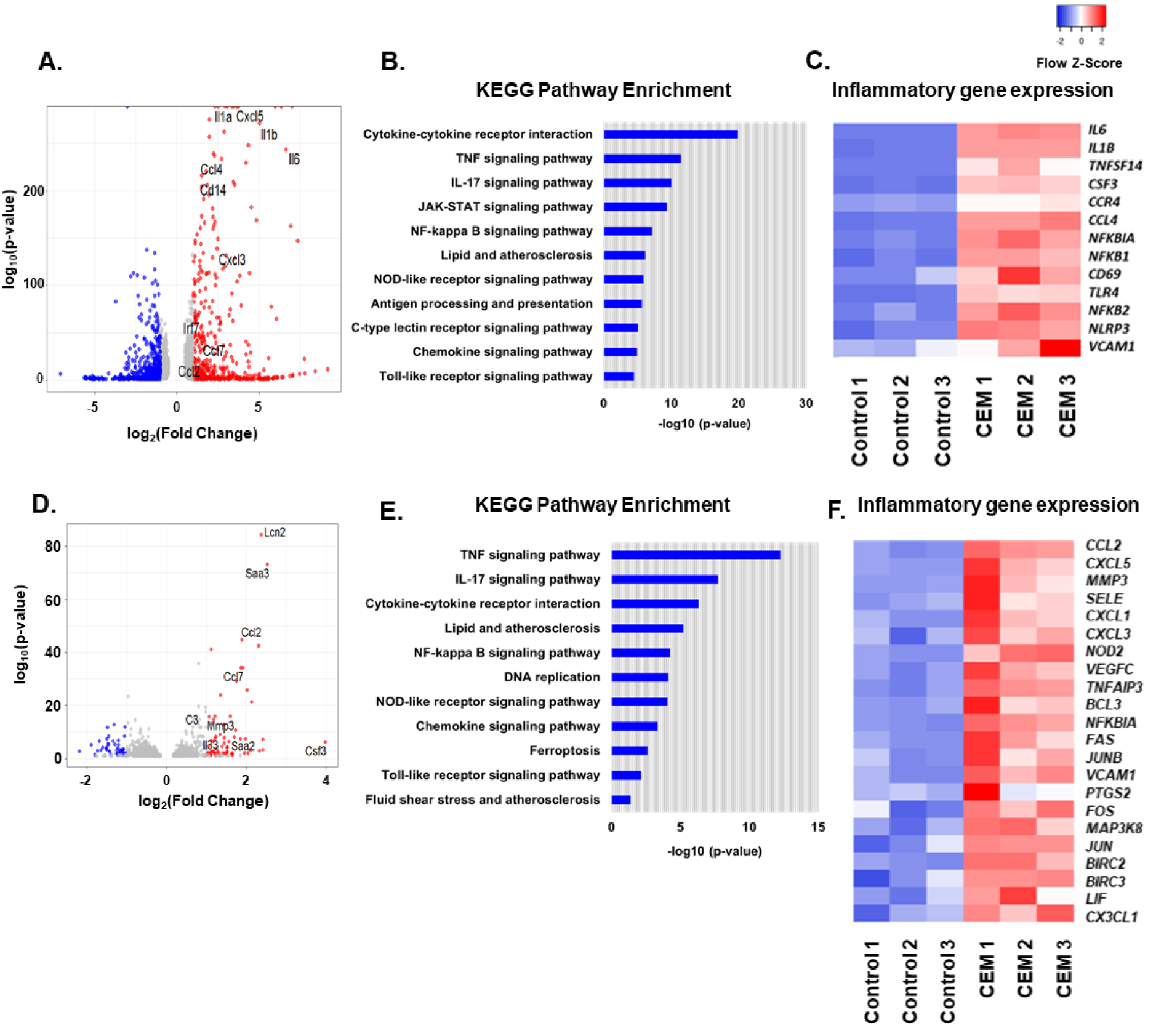
Differential expression analysis between CEM and control medium-treated PMACs or naïve ECs. **A**. Volcano plot illustrating significantly altered gene expression in CEM versus control medium-treated PMACs, determined by DESeq2 analysis with threshold values (absolute log2 fold change > 1, adjusted p < 0.05). Dashed lines and shaded regions indicate these thresholds, with red showing upregulated genes, blue indicating downregulated genes, and gray showing non-significant changes. Adjusted p-values are shown on a –log10 scale. **B.** KEGG pathway enrichment analysis of the upregulated genes in CEM-treated PMACs (p < 0.05). KEGG pathways were ranked by statistical significance, with the bar length representing the adjusted p-value score on a –log10 scale. Longer bars indicate higher significance. **C.** Heatmap of genes highly enriched in inflammation. Hierarchical clustering was performed using Euclidean distance and average linkage. Colors represent z-scored expression values, with red indicating higher expression and blue indicating lower expression. **D.** Volcano plot illustrating significantly altered gene expression in CEM versus control medium-treated naïve ECs, determined by DESeq2 analysis with threshold values (absolutelog2 fold change > 1, adjusted p < 0.05). Dashed lines and shaded regions indicate these thresholds, with red showing upregulated genes, blue indicating downregulated genes, and gray showing non-significant changes. Adjusted p-values are shown on a –log10 scale. **E**. Heatmap of CEM-treated naïve EC gene signatures related to inflammation. Hierarchical clustering was performed using Euclidean distance and average linkage; colors represent z-scored expression values. (F) KEGG pathway enrichment analysis of CEM-treated naïve EC gene signatures (p < 0.05).

Inflammatory signals have been shown to lead to diminished fatty acid oxidation and increased TG synthesis in macrophages ^39,40^. Consistent with this, the pro-inflammatory gene expression profile was associated with downregulation of genes associated with fatty acid oxidation, such as *Ucp2, Acot1*, and *Acot2*. Genes associated with *de novo* lipid synthesis pathways (such as *Fasn*) were downregulated, suggesting that CEM EVs constitute a source of lipids for macrophage LD accumulation (**Fig. 5C, S.Table1**). These results indicate that CEM triggers remodeling of lipid metabolism and activation of innate immunity and inflammation pathways in macrophages.

### CEM alters gene expression in ECs

We next assessed whether treatment with CEM would have a similar effect on naïve ECs. As observed in macrophages, treatment with CEM led to the upregulation of the same inflammatory pathways (TNF, IL17 and NFkB signaling pathways, and cytokine-cytokine receptor interaction) in naïve ECs (**Fig. 5D-F, S. Table 3 and 4**). This suggests that EV-mediated cross-talk between ECs can promote further inflammation upon chylomicron uptake through an autocrine EC mechanism. Moreover, the upregulation of ICAM and VCAM imply that the treated ECs are now poised to attract more circulating monocytes and drive vascular inflammation.

### EVs from chylomicron-treated ECs contain distinct miRs

Next-generation miR sequencing and differential expression analysis indicated that 31 miRs were downregulated, and 27 miRs were elevated in CEM EVs as compared to those from control media. These changes are shown in the volcano plot with a log2 fold change threshold of |1.0| and a p-value cutoff of < 0.05 (**Fig. 6A**). Although the number of differentially expressed miRs in secreted EVs was considerably lower than the number of transcriptional changes noticed directly in ECs, our analysis allowed for the identification of statistically significant miR signatures. The expression of the top 20 significantly upregulated and downregulated miRs, along with their consistent changes across samples, are depicted in the heatmaps (**Fig. 6B, 6C**). Furthermore, we predicted target genes for specific miRs using miR target filter module implemented in QIAGEN’s Ingenuity Pathway Analysis (IPA) tool.

**Figure 6:**
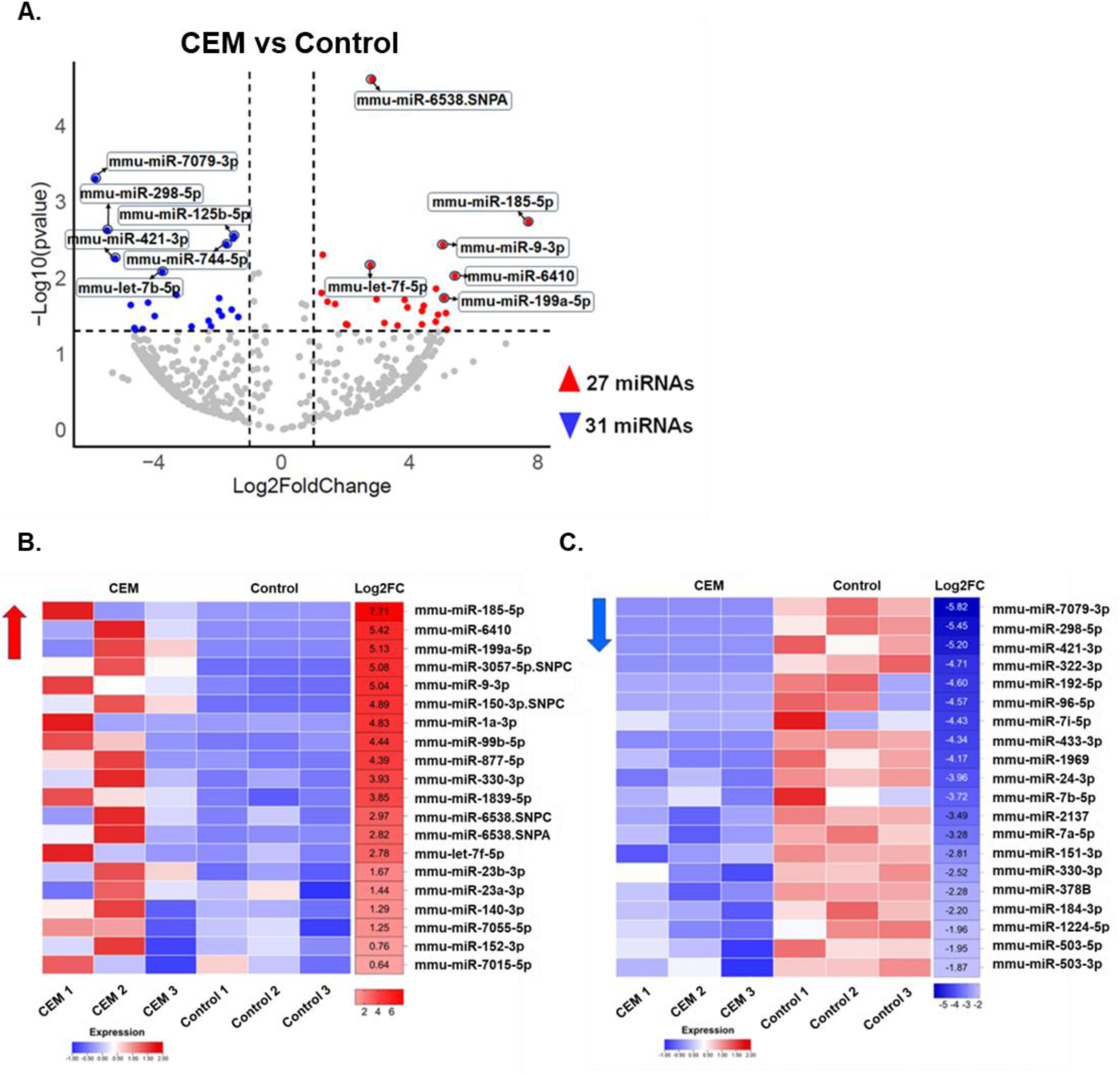
Differential Expression of miRs in CEM vs control conditioned media. **A.** Volcano plot illustrating significantly differentially expressed miRs, determined using a threshold of |log2 fold change| ≥ 1.0 and p < 0.05. Heatmap of the top 20 upregulated miRs (**B**) and top 20 downregulated miRs (**C**), highlighting consistent expression across samples. The color scale represents log2 fold change values, with red indicating higher expression and blue indicating lower expression.

The IPA analysis revealed several upregulated and downregulated miRs that are involved in the control of critical cellular and inflammatory pathways. The downregulated miR mmu-let-7i-5p showed a significant decrease with log2 fold change of −4.43. It targets and negatively regulates a diverse set of genes, including *CCL22*, *CCR7*, *Defa3*, *IFNA4*, *IL10*, *IL12A*, *IL13*, *TRIM6* and *AQP4*, which are predominantly involved in inflammatory response involvinginterleukin, macrophage alternative activation, and cytokine signalling pathways ^41^. Another significantly downregulated miR, mmu-miR-24-3p (with log2 fold change of −3.96) targets and negatively regulates the gene TLR8, implicated in activation of innate immune signalling ^42^. The miR mmu-miR-199a-5p (upregulated with a log2 fold change of 5.13) inhibits the gene *IRF4*, an inhibitor of TLR that blocks production of proinflammatory cytokines in response to TLR stimulation ^43,44^. Hence, endothelial chylomicron uptake appears to induce release of pro-inflammatory miRs. Other changed miRs included mmu-miR-1a-3p (increased with a log2 fold change of 4.48), which inhibits the gene *CAV2* essential for caveolar function and EC signalling, and influences vascular remodelling and responses to shear stress ^45^. These results show that chylomicron treatment modifies the profiles of EC miR release, which may influence function of ECs and macrophages, and induce some of the dysfunctional responses linked to increased chylomicron levels ^46^.

### EC SR-BI knockdown reduces the inflammatory effect of EVs

Chylomicron uptake by ECs was inhibited by knocking down the SR-BI receptor with an antisense oligonucleotide (ASO) ^14^. Surprisingly, the number of EVs released by chylomicron-treated ECs was not reduced by SR-BI knockdown (**Fig. 7**). However, the inflammatory effects of those EVs on underlying ECs were attenuated in cells treated with CEM from SR-BI knockdown ECs (**Fig. 7A and B, S. Table 3**). For instance, genes that are hallmarks of leucocyte recruitment and inflammation such as *Ccl2, Cxcl1* and *Cxcl5* were less expressed in ECs treated with CEM from SR-BI knockdown ECs. In addition, we found that the miR cargo of the secreted EVs differs with the knockdown of SR-BI. Thus, blocking chylomicron uptake from ECs can impair the inflammatory response intracellularly and through miR content of EVs.

**Figure 7.**
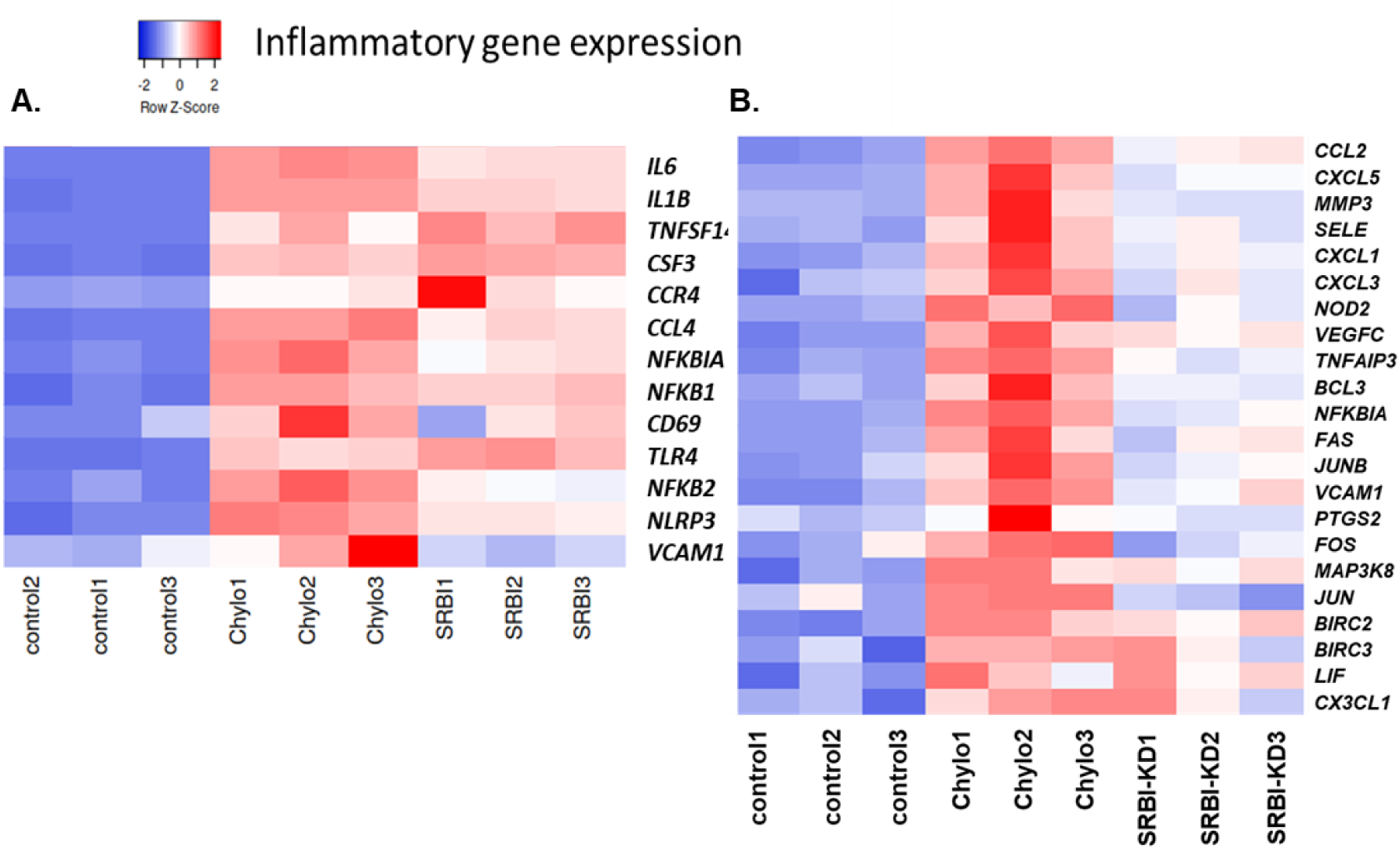
Differential gene expression analysis of naïve ECs treated with CEM from control or SR-BI knockdown ECs, highlighting attenuated inflammation. (A–B) Heatmaps of differentially expressed genes, identified using DESeq2 analysis (absolute log2 fold change > 1, adjusted p < 0.05), that are highly enriched in immune-related KEGG pathways and inflammatory signatures. Hierarchical clustering was performed using Euclidean distance and average linkage. Colors represent z-scored expression values, with red indicating higher expression and blue indicating lower expression.

## DISCUSSION

ECs are a major source of circulating EVs ^17–20^, which likely mediate their communication with parenchymal cells. Crewe et al. previously showed that caveolin 1 is transferred from ECs to adipocytes via EVs and that this process is modulated by nutrient status (fasting/refeeding) as well as obesity ^20^. We previously showed that ECs secrete small EVs that transfer circulating fatty acids to muscle and these sEVs influence transcription of a number of metabolic genes when added to myotubes ^16^. We have also shown that ECs internalize chylomicrons through an SR-BI-dependent pathway, as likely occurs in the postprandial period ^13,14^.. The internalized chylomicrons undergo lysosomal hydrolysis and some of the released lipid promotes formation of perilipin2-coated LDs ^14^. Here, we show that EC uptake of chylomicrons also leads to secretion of EVs, which contain primarily PLs in addition to proinflammatory cargo including miRs linked to inflammation. These EVs lead to major changes in the genetic repertoire of macrophages and ECs.

We first showed that chylomicron treatment doubled the number of EVs released by ECs. Overall, surface markers and lipid composition of the EVs did not change. We tracked the movement of chylomicron fatty acids using rat chylomicrons labeled with [^13^C]oleic acid. Overwhelmingly after a duodenal infusion, the ^13^C labeled oleate was incorporated into chylomicron TG. Some of this label was retained in ECs, likely within LDs. The released EVs primarily contained ^13^C-labeled PLs and little label appeared in EV TGs or NEFAs;.

This surprising observation led us to carefully analyze the labeled lipids from both the ECs and EVs. To enhance our understanding of fatty acid metabolism and ensure consistent data interpretation across different analytical systems, we focused on the overall quantification of PLs and present our lipidome data interpretation at the class level, which highlights the most important aspect of this analysis, that the chylomicron TGs underwent intracellular lipolysis and the released NEFAs and glycerolipids were used for TG and PL synthesis within ECs, while the secreted label predominantly was recoved in PLs. Thus, the transcellular movement of chylomicron TG led to intracellular lipolysis and rescretion of the acyl chains in PL. This pathway could provide FAs for oxidation by the heart and other tissues that rely on FA oxidation for optimal energy production.

We proved that the CEM has a remarkable effect on the transcriptome of PMACs and naïve ECs. We found that pathways that lead to pro-inflammatory responses and recruitment of leukocytes such as TNF-α, NFK-ß and IL-17 signaling pathways were upregulated in both cell types only when cells were exposed to CEM. At the same time, a cluster of upregulated genes belongs to the “lipid and atherosclerosis” pathway, including metalloproteinases (MMP3, MMP1b), interleukins (e,g., *Il6, Il1b*) and adhesion molecules (e.g., *Vcam1*). Thus, innate immunity in ECs and macrophages is induced by CEM.

EVs are known to be a rich source of miRs. In silico analysis identified several miRs with the potential to alter macrophage metabolism and inflammatory gene expression; on-going research is directed to altering the activity of these miRs. We employed the miR Target Filter in IPA to understand the biological effects of miRs, using experimentally validated interactions from TarBase and miRecords, along with predicted miR-mRNA interactions from TargetScan. We connected the differentially expressed miRs with the transcriptome of the PMACs in order to pinpoint mechanisms of CEM -driven inflammation. The current analysis emphases changes in miR expression profiles in chylomicron-treated ECs, indicating a complex regulatory system that may influence endothelial function via miR mediated pathways. The observed downregulated miRs such as mmu-let-7i-5p and mmu-miR-24-3p, as well as the overexpression of others such as mmu-miR-1a-3p and mmu-miR-199a-5p, highlights a complex interplay of gene expression modulation that may be critical for cellular responses to lipid load.

ASO-knockdown of SR-BI expression led to several changes in chylomicron-treated EC release of EVs, but did not reduce the overall numbers of EVs in the media. However, EVs from SR-BI knockdown ECs decreased induction of inflammatory pathways in ECs and macrophages. These changes may be due to differences in miR cargo as a result of reduced chylomicron uptake in SR-BI KD ECs.

What is the *in vivo* relevance of our studies? Aside from lipolysis, the major route for delivery of chylomicron lipids to parenchymal cells, we show a second pathway whereby dietary lipids alter cellular function. EC uptake of chylomicrons alters their production of EVs that can change the biology of underlying macrophages, as well as other ECs. LDs accumulate in the aorta during the postprandial period ^13^ and that this process is exaggerated during the hyperchylomicronemia associated with LpL deficiency ^14^. EC LDs appear to be part of a novel transEC lipid transport mechanism in which acyl groups in TG are converted to EV PLs. In arteries that are largely devoid of LpL ^47^, the changes in ECs suggest a process that creates postprandial EC dysfunction, this may be responsible for the vasodilation defects in humans after a high fat meal ^48,49^. Such effects have been linked to the development of vascular disease ^48^. Postprandial lipemia is a well-established marker for cardiovascular disease development in humans and we postulate that EC uptake of chylomicrons leading to EV production is one pathway exacerbating vascular dysfunction and atherosclerosis development. Not only does EC chylomicron uptake drive macrophage inflammation but this process can also affect neighboring cells via EC cross-talk.

## ACKNOWLEDGMENTS

This work was supported by grants HL164949, HL151328 and HL160470 to IG, K08-DK117065 and P01-HL-160470-5346 to JOA, HL160646 to EDL and an American Heart Association postdoctoral fellowship award to AGC. We acknowledge the support of the NYU Research Resource Identifier number RRID:SCR_022444.

## TABLES

**Supplement Table 1.** List of Differentially Expressed Genes (DEGs) in Chylomicron treated vs. control ECs.

**Sheet 1:** DEGs_ECs

**Sheet 2:** KEGG pathways_ECs

**Supplement Table 2**. List of Differentially Expressed Genes (DEGs) in CEM vs. control medium-treated PMACs.

**Sheet 1:** DEGs_PMACs

**Sheet 2:** KEGG pathways_PMACs

**Supplement Table 3**. List of Differentially Expressed Genes (DEGs) in CEM vs. control medium-treated naïve ECs.

**Sheet 1:** DEGs_naiveECs

**Sheet 2:** KEGG pathways_naiveECs

**Supplement Table 4**. List of Differentially Expressed Genes (DEGs) in Chylomicron treated EC SR-BI knockdown vs. control ECs.

**Sheet 1:** DEGs_ EC SR-BI_KD_vs_control

**Supplement Figure 1. <u>IPA canonical pathway “Atherosclerosis Signaling” in</u> <u>PMAC gene expression analysis.</u>** This pathway illustrates known gene networks of atherosclerosis development among endothelial cells, macrophages, and smooth muscle cells. Nodes represent genes, molecules, or complexes in this pathway, and lines/arrows between nodes indicate known relationships from the Ingenuity Knowledge Base. Nodes outlined in purple indicate molecules that were measured as differentially expressed in our dataset, with the intensity of the colored fill reflecting the level of up- (red) or down- (green) regulation. The blue- and orange-colored molecules and lines denote predicted activation states from the Molecular Activity Predictor function in Ingenuity Pathway Analysis (IPA): blue indicates predicted inhibition, and orange indicates predicted activation. Yellow lines indicate relationships where our findings are inconsistent with the predicted state of the downstream molecule. Broad lines with explanatory text beside the pathway denote the cellular locations of the molecules. Each node’s shape signifies its functional class: Nested Circle/Square = Group/Complex, Horizontal Ellipse = Transcriptional Regulator, Vertical Ellipse = Transmembrane Receptor, Vertical Rhombus = Enzyme, Square = Cytokine/Growth Factor, Triangle = Kinase, Circle = Other. Solid lines represent direct relationships, while dashed lines represent indirect relationships.

